# Rapid generation of pigment free, immobile zebrafish embryos and larvae in any genetic background using CRISPR-Cas9 dgRNPs

**DOI:** 10.1101/2021.04.02.438260

**Authors:** Andrew E. Davis, Daniel Castranova, Brant M. Weinstein

**Author notes:** To whom correspondence should be addressed, Brant M. Weinstein, Section on Vertebrate Organogenesis, Building 6B, Room 4B413, 6 Center Drive, Bethesda, MD 20892.

## Abstract

The ability to carry out high-resolution, high-magnification optical imaging of living animals is one of the most attractive features of the zebrafish as a model organism. However, formation of obscuring pigmentation as development proceeds and difficulties in maintaining sustained immobilization of healthy, living animals remain challenges that limit the application of live imaging. Chemical treatments can be used to suppress pigment formation and movement, but these treatments can lead to developmental defects. Genetic mutants can also be used to eliminate pigment formation and immobilize animals but maintaining these mutants in lines carrying other combinations of transgenes and mutants is difficult and laborious. Here, we show that CRISPR duplex guide ribonucleoproteins (dgRNPs) targeting the *slc45a2 (albino)* and *chrna1 (nic1)* genes can be used to efficiently suppress pigment formation in and immobilize F0 injected animals. CRISPR dgRNPs can be used to generate pigment-free, immobile zebrafish embryos and larvae in any transgenic and/or mutant-carrying background, greatly facilitating high-resolution imaging and analysis of the many transgenic and mutant lines available in the zebrafish.

## INTRODUCTION

The zebrafish has become one of the most versatile and important model organisms in use today. Its small size, high fecundity, short generation time, and optically clear early development make the fish highly accessible to experimental and genetic manipulation. Most zebrafish genes have a human ortholog ^1^, highlighting the relevance of zebrafish research to understanding human development and disease. The ability to carry out high-resolution optical imaging of structures deep within the intact, developing animal is one of the most highly prized features of the zebrafish model, but several major challenges limit the application of these methods, including the formation of obscuring pigmentation as development proceeds and difficulties in maintaining sustained immobilization of healthy, living animals.

Although zebrafish are relatively transparent during early embryogenesis, onset of pigmentation begins at approximately 24 hours post-fertilization (hpf), progressively obscuring and limiting imaging of different tissues and structures within the developing animal ^2^. The formation of the highly opaque melanin pigment in melanocytes can be suppressed by treatment with 1-phenyl 2-thiourea (PTU) beginning prior to 24 hpf ^3-5^, but PTU is toxic and prolonged treatment leads to developmental defects ^6, 7^. Genetic mutations that interfere with the development of pigment-producing cells provide an alternative to PTU treatment. Pigment-deficient zebrafish including the “Casper” (*roy orbison, nacre* double mutant) line and *slc45a2 (albino)* mutants have been used as a valuable zebrafish models for developmental and adult imaging studies in the zebrafish, including in cancer and transplantation research ^8, 9^. Casper mutant zebrafish lack iridophores and melanophores due to defects in *mpv17* and *mitfa*, respectively, yielding relatively pigment-free fish that still retain opaque pigmentation of the retinal pigment epithelium ^10^. The *slc45a2* gene is involved in melanocyte differentiation and melanin production and zebrafish *slc45a2 (albino)* mutants lack melanin pigmentation in both the retinal pigment epithelium of the eye and in melanocytes throughout the rest of the animal ^11^. Although these mutants greatly facilitate larval and adult optical imaging, it is extremely time consuming and laborious to cross these mutants into and continue to maintain them together with strains containing other combinations of mutants and transgenes, severely limiting their use in zebrafish imaging experiments.

In addition to optical clarity, high-magnification, high-resolution light microscopic imaging of living zebrafish also requires rigid immobilization and complete suppression of skeletal muscle contraction. Like inhibition of pigment development, immobilization can also be achieved through both chemical and genetic means. Solubilized MS-222 or tricaine methane sulphonate (3-aminobenzoic acid-ethyl ester methanesulfonate) is widely used to anesthetize zebrafish for laboratory research ^4^, but it can have adverse physiological consequences on zebrafish larvae and adults ^12-14^. Zebrafish can also be “genetically immobilized” by knocking out genes coding for proteins involved in skeletal muscle function. The *cholinergic receptor alpha 1* (*chrna1*, aka *nic1*) gene is expressed in skeletal muscle and required for synaptic transmission from motor neurons to muscle cells ^15, 16^. Mutants for *chrna1* exhibit complete skeletal muscular paralysis, although they have normal heartbeat and blood circulation and develop morphologically quite normally for many days. However, as noted above for the pigment mutants, *chrna1* mutants must be crossed into and maintained in whatever genetic background is being imaged, requiring extensive time, effort, and fish tank space. Generating and maintaining lines for obtaining optically clear, immobilized animals carrying specific transgenes and mutants for imaging requires complicated and laborious genetic crosses with very low yields of the appropriate progeny needed for imaging (see Discussion, below).

The advent of high-efficiency targeted CRISPR mutagenesis using dgRNPs ^17^ provides a useful, efficient alternative to toxic chemicals and complex genetic crosses for generating optically clear, immobilized animals for imaging. CRISPR technology has revolutionized reverse genetics in the zebrafish, making it simple and straightforward to generate genetic mutants in virtually any gene of interest. Until recently, the relatively modest efficiency of available CRISPR technologies and concerns about off-target effects in injected animals made it necessary to raise injected animals and then cross their progeny to identify germline carriers for phenotypic analysis of defects in particular targeted genes. However, recent studies have demonstrated significant improvement in targeted CRISPR-Cas9-mediated mutagenesis through the use of a two-RNA component (crRNA and tracrRNA) CRISPR system complexed with Cas9 protein ^17, 18^. When these duplex guide ribonucleoproteins (dgRNPs) are injected into zebrafish eggs, targeted cutting proceeds at a rate that leaves nearly every cell harboring a bi-allelic mutation ^19^. The high level of efficiency and specificity of these new methods makes it possible to generate F0-injected animals that are close to homozygous null for a given gene of interest.

In this study, we show how these new methods can be harnessed to facilitate generation of optically clear, pigment-free and immobile zebrafish embryos and larvae in any transgenic and/or mutant-carrying background. This important new capability will greatly facilitate high-resolution imaging and analysis of zebrafish transgenic and mutant lines, eliminating the need to incorporate pigmentation and paralysis mutants in these lines or the use of chemical treatments.

## RESULTS

We utilized the highly efficient mutagenic activity of dgRNP-Cas9 CRISPR complexes (“dgRNPs”) to target exons 1 and 2 of the *slc45a2 (albino)* gene, required for melanocyte differentiation and melanin pigmentation (**Fig. 1A**), exon 2 of *chrna1 (nic1)*, a gene required for skeletal muscle movement (**Fig. 1B**), and exon 1 of *plxnd1 (obd)*, a gene required in the endothelium for proper vascular patterning (**Fig. 1C**). Fragment size analysis of fluorescent PCR sequences amplified from DNA targeted by each of the dgRNP sequences showed that in contrast to control uninjected animals, DNA from animals injected with dgRNPs targeting *slc45a2* exon 1 (**Fig. 1D,E**), *slc45a2* exon 2 (**Fig. 1F,G**), *chrna1* exon 2 (**Fig. 1H,I**), or *plxnd1* exon 1 (**Fig. 1J,K**) showed a very high proportion of loss of the wild type alleles, as indicated by the presence of a large variety of insertions and deletions (**Fig. 1L**).

**Figure 1.**
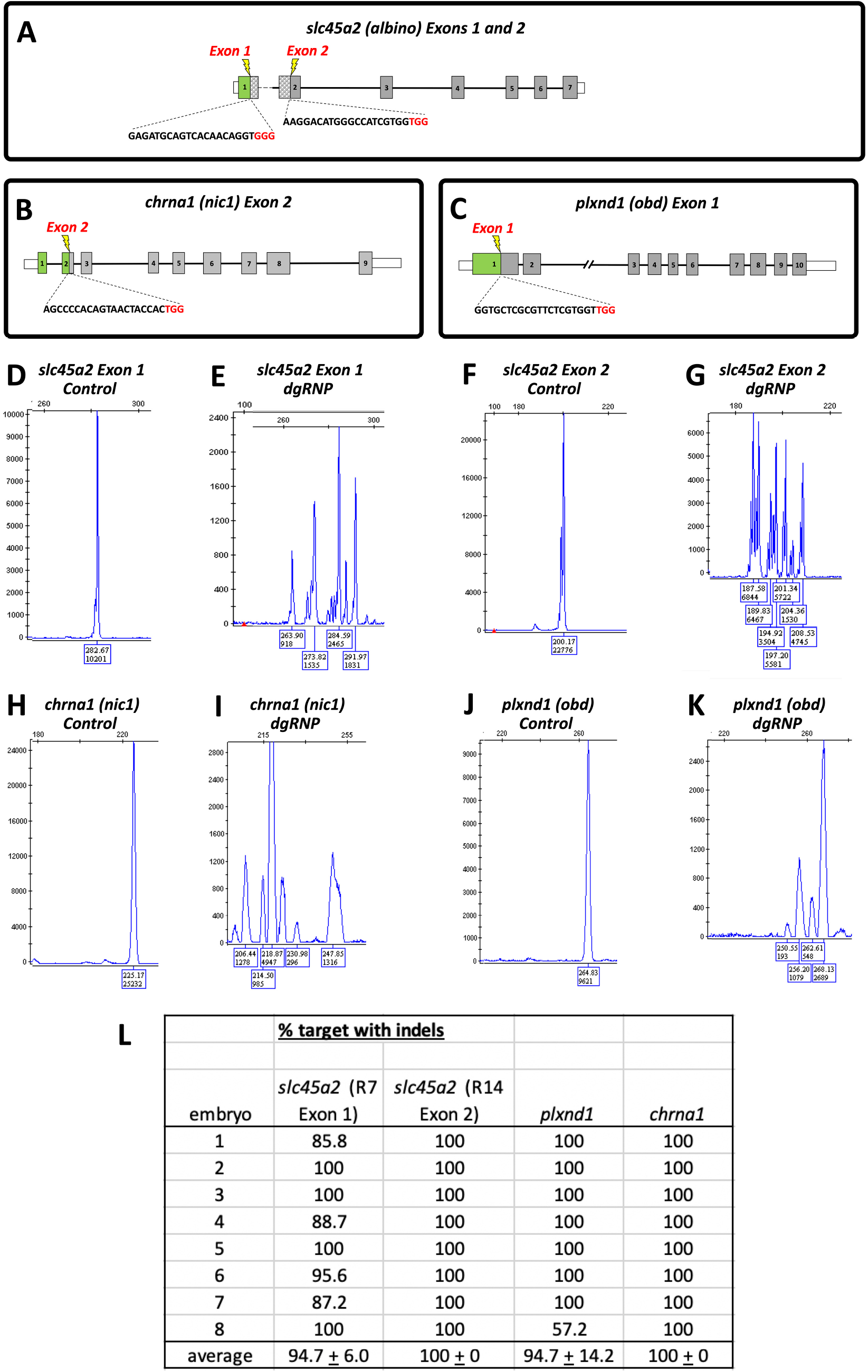
High efficiency targeting of *slc45a2, chrna1*, and *plxnd1* with CRISPR dgRNPs. **A-C**. Schematic diagrams of the (A) *slc45a2 (albino)*, (B) *chrna1 (nic1)*, and (C) *plxnd1 (obd)* loci showing dgRNP guide sequences targeting exons 1 and exon 2 of *slc45a2*, exon2 of *chrna1*, and exon 1 of *plxnd1*. **D,E**. Fragment analysis traces of *slc45a2* exon 1 sequences PCR amplified from 2 dpf control (B) or *scl45a2* exon 1 (C) dgRNP injected animals. **F,G**. Fragment analysis traces of *slc45a2* exon 2 sequences PCR amplified from 2 dpf control (D) or *scl45a2* exon 2 (E) dgRNP injected animals. **H,I**. Fragment analysis traces of *chrna1 (nic1)* exon 2 sequences PCR amplified from 2 dpf control (H) or *chrna1* exon 2 (I) dgRNP injected animals. **J,K**. Fragment analysis traces of *plxnd1 (obd)* exon 1 sequences PCR amplified from 2 dpf control (J) or *plxnd1* exon 1 (K) dgRNP injected animals. **L**. Quantitation of the percentage of PCR amplified target sequences with insertions or deletions (indels) in 8 embryos injected with dgRNPs directed against *slc45a2* exon 1, *slc45a2* exon 2, *chrna1* exon 2, or *plxnd1* exon 1, normalized against fragment analysis traces from 8 control embryos.

Wild type zebrafish embryos and larvae have highly opaque melanin-containing pigment cells that interfere with optical imaging of a variety of different areas of the animal (**Fig. 2A,C**), including the dorsal head and trunk (small arrows in Fig. 1C), eyes (arrowheads in Fig. 1C), ventral trunk (asterisks in Fig. 1C), and yolk cell (large arrows in Fig. 1C). By comparison, a large majority of animals (40/48, 83%) co-injected with *slc45a2* exon 1 and 2 dgRNPs displayed virtually complete loss of melanin pigment (<10 very small pigment cells), including the melanocytes obscuring the dorsal head, dorsal and ventral trunk, pigment epithelium of the eye, and yolk cell (**Fig. 2B,D, Movie 1**). Although injection of dgRNPs targeting *slc45a2* exon 2 alone eliminated most melanin pigment, co-injection of the slightly less efficient *slc45a2* exon 1 dgRNPs (see **Fig. 1L**) resulted in nearly total loss of melanin pigmentation from injected animals (**Fig. 2B,D, Movie 1**). Therefore, a combination of *slc45a2* exon 1 and *slc45a2* exon 2 dgRNPs was used for all subsequent experiments.

**Figure 2.**
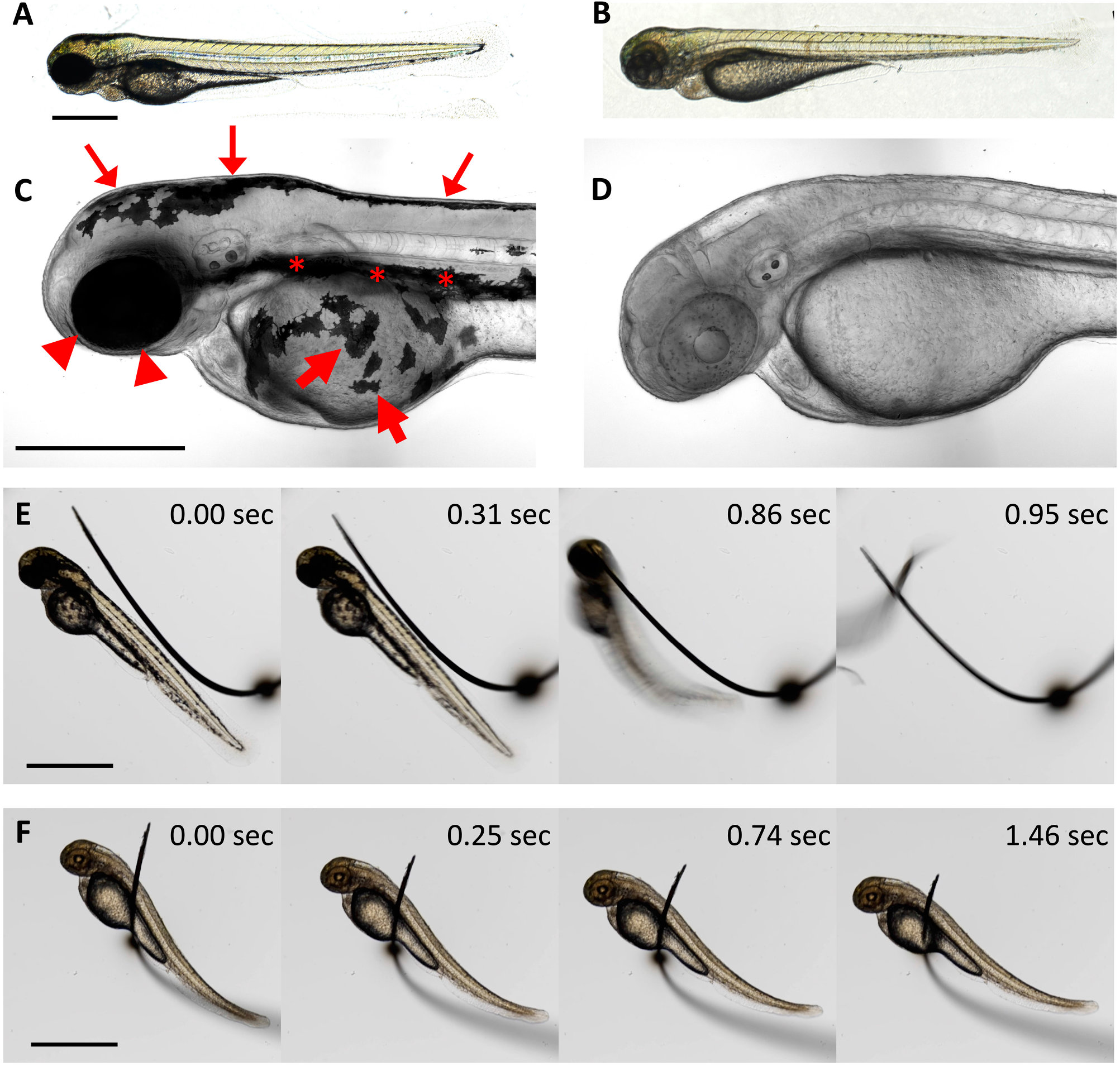
Loss of pigment and paralysis in animals injected with a cocktail of dgRNPs targeting *slc45a2, chrna1*, and *plxnd1*. A. Lateral view of a 2 dpf uninjected control animal. B. Lateral view of a 2 dpf animal injected with a mixture of 4 dgRNPs targeting *slc45a2* exon 1, *slc45a2* exon 2, *chrna1* exon 2, and *plxnd1* exon 1 (“4-dgRNP” animals). C. Lateral view of the head and rostral trunk of a 2 dpf uninjected control animal with clearly evident melanocytes. D. Lateral view of the head and rostral trunk of a 2 dpf 4-dgRNP animal lacking melanin pigment. E. Time series showing touch responsiveness of a 2 dpf uninjected control animal (see Movie 1). F. Time series showing lack of touch responsiveness of a 2 dpf 4-dgRNP animal (see Movie 1). Scale bars = 500 µm (A - D), 1 mm (E,F). Panels E and F show close-ups from selected frames of **Movie 1**.

Wild type zebrafish embryos and larvae also display robust responses to touch and spontaneous movements that make it necessary to either anesthetize the animals with chemical agents or carry out complex genetic crosses to generate animals for imaging that carry “paralyzing” homozygous mutations in genes such as *chrna1*. As expected, control uninjected animals rapidly initiated an “escape response” after being touched or shaken (**Fig. 2E, Movie 1**), and depending on their age underwent occasional spontaneous movements in the absence of stimulation. In contrast, a large majority of animals (45/48, 94%) injected with *chrna1* exon 2 dgRNPs displayed complete loss of both sensitivity to touch and elimination of spontaneous movements (**Fig. 2E,F, Movie 1**). Unlike control animals *chrna1* exon 2 dgRNP-injected animals remained entirely motionless while being manipulated and embedded in mounting media (eg., or agar, methylcellulose) for imaging and during actual imaging. Other than their pigment or paralysis phenotypes, the gross morphology of dgRNP-injected animals was indistinguishable from their control uninjected siblings, no significant developmental delays were noted, and the heartbeat and circulation remained strong.

Having demonstrated that dgRNP targeting of *slc45a2* or *chrna1* could be used to efficiently generate melanin pigment-free or paralyzed embryos and larvae, we examined whether these could be combined together with each other and with dgRNPs targeting other genes to rapidly generate pigment-free, motionless animals for high-resolution imaging the phenotypic consequences of loss of specific genes in any transgenic background of interest.

We injected dgRNPs targeting *slc45a2* exon 1, *slc45a2* exon 2, *chrna1* exon 2, and *plxnd1* exon 1 into *Tg(kdrla:egfp)*^*la116/la116*^, *Tg(gata1:dsred)*^*sd2/sd2*^ homozygous double-transgenic embryos and carried out high-resolution confocal imaging of blood vessels (kdrl:gfp-positive) and blood cells (gata1:dsred-positive) *in vivo* (**Fig. 3A-H, Movie 1**). Quadruple-dgRNP injected (“4-dgRNP”) *Tg(kdrla:egfp)*^*la116/la116*^, *Tg(gata1:dsred)*^*sd2/sd2*^ double-transgenic animals were completely immobilized and displayed a near total lack of pigment that greatly facilitated high-resolution, high-magnification confocal imaging of vessels that were obscured in their uninjected siblings (**Fig. 3C-H**). Vessels such as the dorsal longitudinal anastomotic vessel (DLAV) in the dorsal trunk and the supraintestinal artery in the ventral trunk that were completely or almost completely concealed by melanocytes in control injected animals (asterisks and hash marks in **Fig. 3C,E**) were readily visualized in their 4-dgRNP injected siblings (**Fig. 3D,F, Movie 1**). Imaging of blood vessels in the dorsal head and eyes of uninjected control animals was also confounded by the large number of highly opaque melanocytes on the dorsal surface of the head (asterisks in **Fig. 3G**) and the highly opaque pigment epithelium around the eyes (hash mark in **Fig. 3G**). These vessels were also readily visualized and imaged in their 4-dgRNP injected siblings (**Fig. 3H, Movie 1**). In addition to their immobility and loss of pigment, 4-dgRNP injected animals also showed a robust *“out of bounds”* vascular phenotype comparable to that described in animals homozygous for germline genetic mutations in *plxnd1* ^20^. A large majority of 4-dgRNP injected animals (45/48, 94%) displayed highly disorganized and mispatterned blood vessels, including the intersegmental vessels in the trunk, subintestinal vessels over the yolk cell, and intracranial vessels in the head, strongly reminiscent of the vessel defects previously reported in *plxnd1* homozygous null mutant animals (**Fig. 3B,B,F,H, Movie 1**).

**Figure 3.**
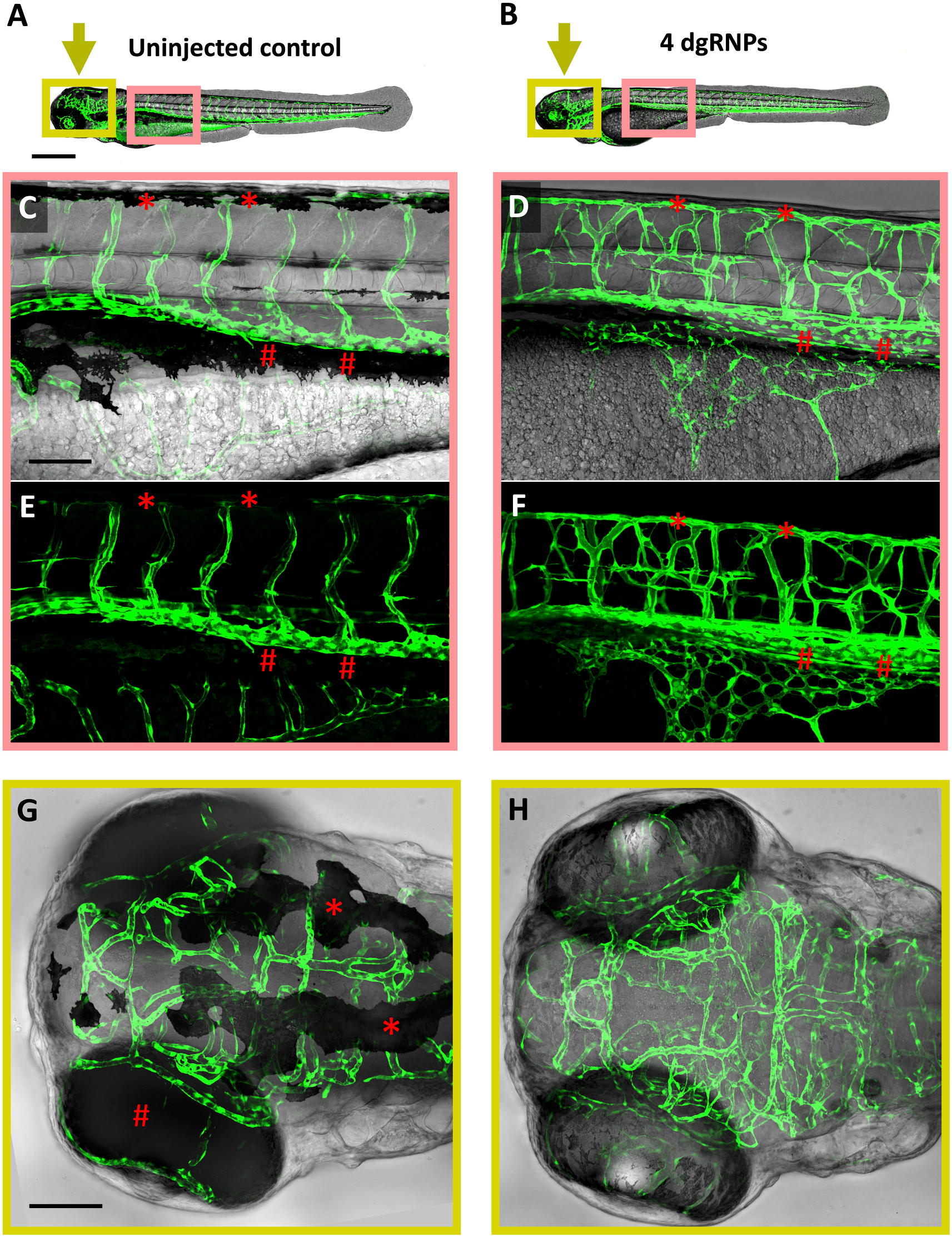
Visualization of the *“out of bounds”* vascular phenotype in 4-dgRNP-injected double-transgenic animals. Loss of pigment and paralysis in 2 dpf *Tg(kdrla:egfp)*^*la116/la116*^, *Tg(gata1:dsred)*^*sd2/sd2*^ double transgenic animals that were either uninjected or injected with a cocktail of four dgRNPs targeting *slc45a2* exon 1, *slc45a2* exon 2, *chrna1* exon 2, and *plxnd1* exon 1 (“4-dgRNP” animals). **A,B**. Confocal (EGFP) and transmitted light composite lateral view images of uninjected control (A) and 4-dgRNP-injected (B) animals. Pink and yellow boxes indicate the approximate locations of the higher magnification images shown in panels C-F and G,H, respectively. Yellow arrows indicate the direction if imaging (dorsal) for panels G,H. **C-F**. Confocal (EGFP) and transmitted light composite (C,D) or confocal (EGFP) only (E,F) higher magnification lateral view images of the mid-trunks of uninjected control (C,E) and 4-dgRNP-injected (D,F) animals. In panels C and E, red asterisks indicate pigment obscuring imaging of the dorsal longitudinal anastomotic vessel, while red hash marks indicate pigment obscuring imaging of the cardinal vein and supraintestinal artery. **G,H**. Confocal (EGFP) and transmitted light composite higher magnification dorsal view images of the heads of uninjected control (C,E) and 4-dgRNP-injected (D,F) animals. In panel G, red asterisks indicate obscuring pigment on the dorsal surface of the head, while the red hash mark indicate obscuring pigment around the eye. Scale bars = 500 µm (A,B), 100 µm (C-H).

## DISCUSSION

In this study we show that by utilizing recent advances in CRISPR technology immobilized, pigment-deficient dgRNP-injected animals can be easily generated in any genetic background, bypassing the complex breeding schemes that would be needed to generate the appropriate homozygous null mutant combinations or the toxic effects of using PTU and tricaine to inhibit pigment formation and movement, respectively. By injecting dgRNPs targeting the *slc45a2 (albino)* and *chrna1 (nic1)* genes into transgenic animals we can obtain a high percentage of pigmentless and immobile embryos usable for extended confocal imaging. This provides a time-and labor-saving alternative to maintaining the *slc45a2* and *chrna1* mutants in multiple lines used for imaging, since injection of *slc45a2* and *chrna1* can be used be used together with any mutant- and/or transgene-carrying line.

Furthermore, even where lines are available carrying all of the necessary transgenes and genetic mutants, use of dgRNPs to inhibit pigment formation and movement provides a huge advantage for imaging experiments. In the example described above where a 4-dgRNP cocktail targeting slc45a2, chrna1, and plxnd1 was injected into *Tg(kdrla:egfp)*^*la116/la116*^, *Tg(gata1:dsred)*^*sd2/sd2*^ homozygous double-transgenic animals, given near 100% cutting efficiency almost every injected animal is paralyzed and virtually pigment-free with a robust out of bounds phenotype (**Fig. 3**, Movie 1). In contrast, even assuming the best-case scenario of starting with an incross of *slc45a2*^*+/-*^, *chrna1*^*+/*^, *plxnd1*^*+/-*^, *Tg(kdrla:egfp)*^*la116/la116*^, *Tg(gata1:dsred)*^*sd2/sd2*^ adults, only approximately 1 in 64 of their progeny would be pigment free, immobile, and displaying the out of bounds mutant phenotype. Generating and maintaining the triple-mutant-, homozygous double-transgenic-carrying line needed to generate this “best-case” cross would be a difficult and time-consuming task requiring a substantial commitment of tank space and labor for multiple generations even to maintain this one line, not to mention if multiple lines need to be maintained in this manner.

Although treatment with PTU and tricaine provides an alternative method for generating pigment-free, immobilized animals for imaging, extended treatment with either or both of these chemicals has been shown to lead to developmental and circulatory function defects ^7, 13^. This is of particular concern when carrying out time-lapse imaging of vascular development in the fish. Although mild general developmental defects were noted in a small proportion of animals injected with the 4-dgRNP cocktail, most animals were indistinguishable from their uninjected siblings and morphologically “normal” animals with strong phenotypes could easily be selected for imaging. Like homozygous *slc45a2* mutant animals, 4-dgRNP-injected animals maintain a strong heartbeat and robust circulatory flow for many days. In contrast, animals maintained in tricaine for a day or more invariably display stunted growth, weakened heartbeat, and reduced circulatory flow, making long-term imaging of vascular development in tricaine-treated animals particularly problematic. Although we reported the results of using 4-dgRNP injected animals here to demonstrate the utility of dgRNPs for directly generating “imaging-ready” animals with loss-of-function phenotypes in specific targeted genes of interest, injection of a 3-dgRNP cocktail targeting *slc45a2* exon 1, *slc45a2* exon 2, and *chrna1* exon 2 could also be used to generate “imaging-ready” animals from incrosses of adults carrying heterozygous germline mutations in the same genes of interest, with 1 in 4 animals showing the phenotype of loss of the gene. The development of 3-dgRNP injected animals is indistinguishable from their uninjected siblings (data not shown), and in this case the wild type and mutant animals being compared are both injected with the same 3-dgRNP cocktail.

The profusion of available transgenic lines and mutants in the zebrafish provides tremendous opportunities for extended high-resolution microscopic analysis and imaging of many different developmental events and processes. However, bringing these together in a form amenable to long-term imaging of diverse structures can be very challenging. Using dgRNPs to generate pigment free, immobilized animals provides a powerful new tool for generating animals for imaging on a far shorter timescale with the use of far fewer animals than relying on crosses with germline mutants to incorporate these features.

## MATERIALS AND METHODS

### Zebrafish Methods and Transgenic lines

Fish were housed in a large zebrafish dedicated recirculating aquaculture facility (4 separate 22,000L systems) in 6L and 1.8L tanks. Fry were fed rotifers and adults were fed Gemma Micro 300 (Skretting) once per day. Water quality parameters were routinely measured and appropriate measures were taken to maintain water quality stability (water quality data available upon request). Zebrafish husbandry and research protocols were reviewed and approved by the NICHD Animal Care and Use Committee. CRISPR/Cas9 injections were carried out in the previously published *Tg(kdrla:egfp)*^*la116/la116*^, *Tg(gata1:dsred)*^*sd2/sd2*^ double transgenic line ^21^.

### Imaging and Microscopy

Images were acquired using a Nikon Ti2 inverted microscope with Yokogawa CSU-W1 spinning disk confocal, Hamamatsu Orca Flash 4 v3 camera. Color images were taken using Nikon DS-Ri-2 color camera. The following Nikon objectives: 2X Air 0.1 N.A. and 20X long working distance water immersion 0.95 N.A. were used. Transmitted light images are shown using Extended Depth of Field (EDF) focus stacking (NIS - Nikon Elements AR v5). Embryos were mounted in MatTek glass bottomed 35 mm dishes (P35G-1.5-14-C). for high resolution imaging (20X) embryos were mounted in 0.8% low melting point agarose.

### CRISPR/Cas9 Generation of Zebrafish Mutants

Mutations in the zebrafish *slc45a, chrna1* and *plxnd1* genes were generated using the CRISPR/Cas9 system. gRNA targets and corresponding genotyping primers were chosen from Ensembl GRCz11 using the CHOPCHOP (version 3) web tool ^22^ and are listed below. Target-specific Alt-R® crRNA, universal Alt-R® tracrRNA, Alt-R® S.p. Cas9 nuclease, v.3 and Duplex buffer were purchased from Integrated DNA Technologies (IDT). crRNA:tracrRNA duplex preparation, Cas9 RNP complexing and dgRNP injections were carried out as previously described ^19^.

Guide RNA target sites used in this study:

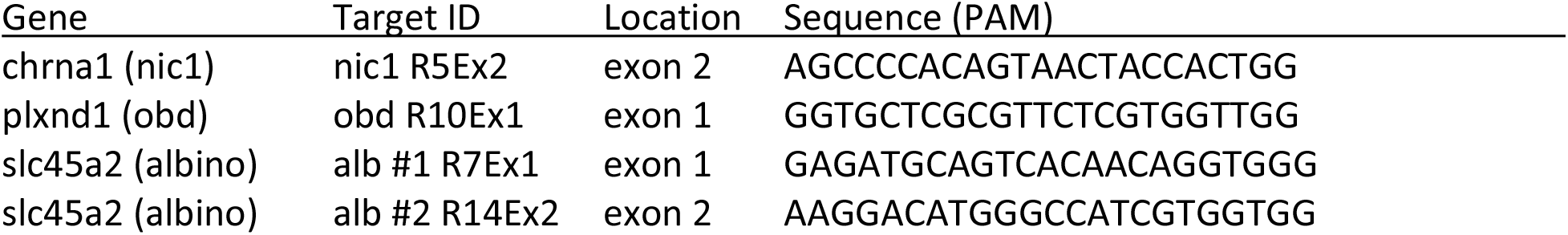

Primer pairs used to detect indel mutations by fluorescent PCR and capillary electrophoresis (M13 / PIG tail sequences):

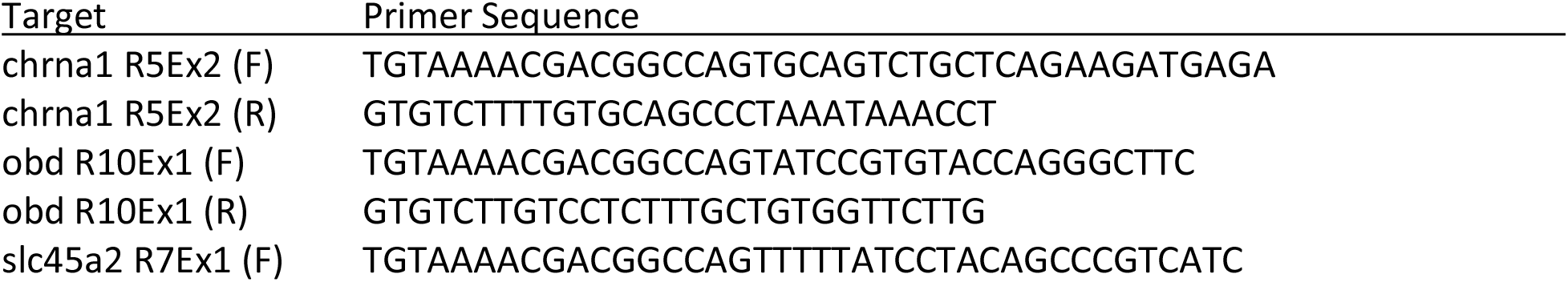
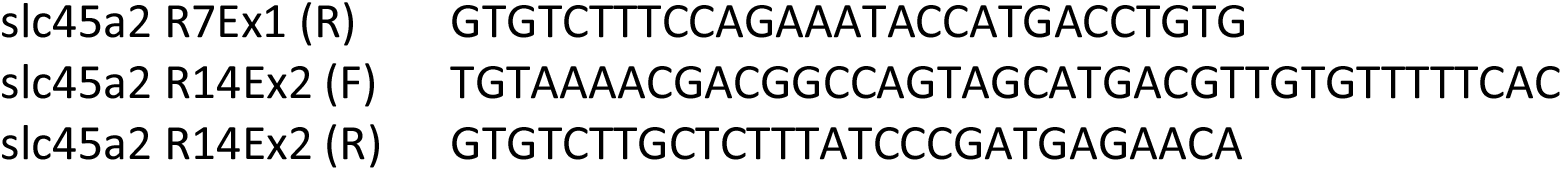

### Fluorescent PCR and DNA Fragment Analysis

The following protocols were modified from previously published methods ^23^. PCR protocol with AmpliTaq Gold DNA Polymerase 1x (10ul) Rxn: 1ul 10x PCR Gold Buffer; 0.5ul MgCl2 25mM; 1ul 0.5mM Fwd primer; 1ul 1mM Rev primer; 0.2ul 10mM FAM-M13 primer; 0.1ul dNTP Master Mix; 0.1ul TaqGold polymerase; 1ul of 1:10 diluted crude gDNA; 5.1ul H20

TaqGold PCR Program: 95°C 10min; 95°C 30sec; 58°C 30sec; 72°C 30sec (1 min/kb); Go To Step 2 x34; 72°C 10min; 15°C Hold. Run on the DNA Analyzer (ABI 3730, Thermo Fisher Scientific) immediately or store at 4°C in the dark for 24 hours max.

3730 Plate set-up: HiDi Formamide/ROX master mix-0.2ul ROX400HD; 9.8ul HiDi Formamide; add 10ul of master mix to each 3730 plate sample well; add 2ul of fluorescent PCR product; cap wells and denature at 95 °C for 5min; uncap all wells and replace with 96 well plate septa to run on the 3730. Follow manufacturer directions to utilize the 3730 DNA Analyzer.

Quantitation of the mutagenic activity of each dgRNP was performed via PCR-amplified DNA fragment trace analysis as previously described ^19^. Raw data from 8 injected and 8 uninjected control embryos was generated by a 3730 DNA Analyzer and uploaded to the Thermo Fisher Connect website. Analysis was performed in the Microsatellite Analysis app where the RFU detection threshold for all FAM-labelled DNA fragments was set at 150 for all samples. Injected embryos where 100 % of analyzed targets had indels had no detectable wild type DNA fragment above threshold levels within the overall pool.

### Study Approval

Zebrafish husbandry and research protocols were reviewed and approved by the NICHD Animal Care and Use Committee at the National Institutes of Health. All animal studies were carried out according to NIH-approved protocols, in compliance with the Guide for the Care and use of Laboratory Animals.

## AUTHOR CONTRIBUTIONS

AED and DC performed experiments; AED, DC, and BMW analyzed results and made the figures; AED, DC and BMW designed the research and wrote the paper.

## ACKNOWLEDGEMENTS

This work was supported by the intramural program of the *Eunice Kennedy Shriver* National Institute of Child Health and Human Development (ZIA-HD001011 and ZIA-HD008808, to BMW).

## FIGURE AND MOVIE LEGENDS

### Movie 1

Pigmentation, paralysis, and vascular phenotypes in 2 dpf *Tg(kdrla:egfp)*^*la116/la116*^, *Tg(gata1:dsred)*^*sd2/sd2*^ double transgenic 4-dgRNP-injected animals. T=0-16” Live imaging of control (right) and 4-dgRNP-injected (left) animals, showing lack of pigmentation and lack of touch responsiveness in the 4-dgRNP-injected animals compared to controls. T=16-22” Higher magnification live image of a 4-dgRNP-injected animal, with box appearing at 18-22” noting the approximate location of higher-magnification transmitted and confocal light images in frames T=22-47”. T=22-47” Higher magnification live transmitted light (T=22-33”) and green/red (egfp/dsred) confocal (T=33-47”) images of a single plane in the trunk of a 4-dgRNP-injected *Tg(kdrla:egfp)*^*la116/la116*^, *Tg(gata1:dsred)*^*sd2/sd2*^ double transgenic animal, showing aberrant “out of bounds” circulation and vascular patterning. T=47-58” Higher magnification green/red (egfp/dsred) confocal image reconstructions of the trunk of a 4-dgRNP-injected *Tg(kdrla:egfp)*^*la116/la116*^, *Tg(gata1:dsred)*^*sd2/sd2*^ double transgenic animal, showing aberrant “out of bounds” vascular patterns. T=58-63” Green/red (egfp/dsred) lateral view confocal image a 4-dgRNP-injected *Tg(kdrla:egfp)*^*la116/la116*^, *Tg(gata1:dsred)*^*sd2/sd2*^ double transgenic animal.

